# Exploring the lower thermal limits for transmission of human malaria, *Plasmodium falciparum*

**DOI:** 10.1101/603787

**Authors:** Jessica L. Waite, Eunho Suh, Penelope A. Lynch, Matthew B. Thomas

## Abstract

The rate of malaria transmission is strongly determined by parasite development time in the mosquito, known as the extrinsic incubation period (EIP), since the quicker parasites develop, the greater the chance that the vector will survive long enough for the parasite to complete development and be transmitted. EIP is known to be temperature dependent but this relationship is surprisingly poorly characterized. There is a single degree-day model for EIP of *Plasmodium falciparum* that derives from a limited number of poorly controlled studies conducted almost a century ago. Here, we show that the established degree-day model greatly underestimates the rate of development of *P. falciparum* in both *Anopheles stephensi* and *An. gambiae* mosquitoes at temperatures in the range of 17-20°C. We also show that realistic daily temperature fluctuation further speeds parasite development. These novel results challenge one of the longest standing models in malaria biology and have potentially important implications for understanding the impacts of climate change.

## 1. Introduction

The transmission of vector-borne diseases is strongly influenced by environmental temperature [1,2]. For this reason, there is considerable interest in the possible effects of climate change on the dynamics and distribution of diseases such as malaria (e.g. [3-6]). One of the key temperature dependencies in malaria transmission is the Extrinsic Incubation Period (EIP; also defined as the duration of sporogony), which describes the time it takes following an infectious blood meal for parasites to develop within a mosquito and become transmissible [7].

Most mechanistic models of *Plasmodium falciparum* transmission base estimates of EIP on the degree-day model of Detinova [8]:

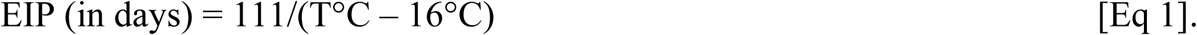

In this model, 111 is the cumulative number of degree-days required for the parasite to complete development once temperature exceeds a lower developmental threshold, T is the average ambient environmental temperature, and 16°C is the lower temperature threshold. However, in spite of widespread use for over 50 years, the Detinova degree-day model is poorly validated (reviewed in [7]). For example, the model was parameterized with limited data from a single study conducted in the 1930’s using the Eurasian vector, *Anopheles maculipennis* [9]. This work provided no empirical measurements of EIP below 20-21°C. Similar historic studies either lacked temperature control, adequate sampling, or did not use *P. falciparum* parasites (e.g. [10-13]). To date, virtually no published studies have measured EIP at cooler temperatures, or confirmed the lower developmental threshold. Further, the model is based on constant temperatures, yet temperatures in the field exhibit diurnal fluctuation, which could affect parasite development [14].

Here we use *An. stephensi* and *An. gambiae* mosquitoes to determine the lower temperature threshold of *P. falciparum*, evaluate EIP at temperatures below 20°C, and examine whether the degree-day model is robust to daily temperature variation.

## 2. Material and methods

### (a) Experimental treatments

Mosquitoes were reared at 27°C following standard protocols [15]. *P. falciparum* (NF54) parasite cultures were either provided by the Johns Hopkins Malaria Institute Core Facility, or produced in our lab following protocols described in [15]. In all cases gametocyte cultures reached approximately 2-4% mature gametocytemia and were between 14-17 days post gametocyte induction when cultures were fed to 3-5 day old mosquitoes. After 20 minutes, blood-fed females were moved to temperature-controlled incubators, set at 80 ± 5% RH. Mosquitoes were sampled over time with midguts and salivary glands dissected to estimate oocyst and sporozoite infection (sampling intervals given in electronic supplementary material, Table S1).

For *An. stephensi* (Liston), a dominant malaria vector in India, Asia, and parts of the Middle East [16], we examined constant temperatures of 16, 17, 18, and 20°C, and fluctuating temperature regimes of 14 ± 5, 16 ± 5 and 18 ± 5°C. For *An. gambiae* (NIH G3), the primary vector in sub-Saharan Africa [17], we examined temperatures of 17, 19 and 19 ± 5°C. The daily fluctuations followed a previously described temperature model [3,14]. Diurnal temperature ranges (DTR) of 5-20°C are common across many malaria transmission settings [3,14,18] and so DTR of 10°C is a representative intermediate value. For each temperature we had 1-6 biological replicates using separate infectious feeds, with at least 150 mosquitoes per infectious feed (see Table S1 for details of replicate numbers and sample sizes). For each feed we included a control set of mosquitoes maintained at 27°C to confirm infections.

### (b) Mosquito survival curves at constant and fluctuating temperature

How EIP affects malaria transmission depends in part on mosquito survival; the absolute duration of EIP does not matter so much as what proportion of mosquitoes survive the EIP to become infectious [15]. Therefore, to parallel the 18°C infection studies, we generated survival curves for adult *An. stephensi.* Mosquitoes 3-5 days old, reared in identical conditions to those in the infection studies, were provided a blood meal and then placed into incubators at either constant 18°C or 18 ± 5°C. Mosquitoes were housed in cups in groups of 10, with a total of 200 mosquitoes per treatment. Cups were examined daily until the last mosquito died.

## 3. Results

The results of the infection experiments are summarized in Table 1 (details of sampling and parasite measures are given in Table S1).

**Table 1.**
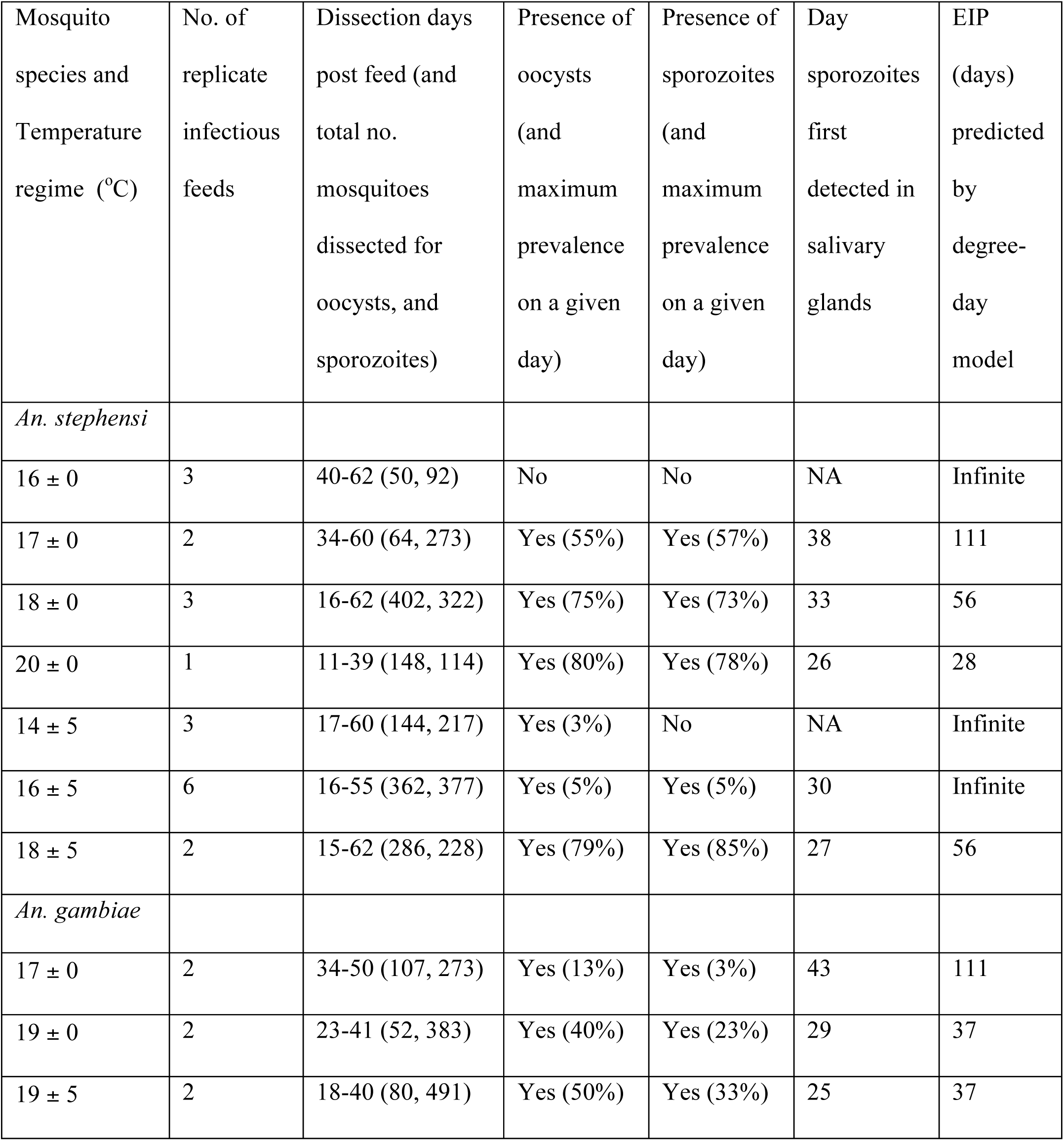
Summary of *P. falciparum* infection across a range of temperatures, showing the number of replicate infectious feeds, the number of mosquitoes dissected to examine for oocysts and sporozoites, the day post-blood-feed when sporozoites were first observed in mosquito salivary glands, and the equivalent EIP predicted by the Detinova degree-day model [8].

| Mosquito species and Temperature regime (°C) | No. of replicate infectious feeds | Dissection days post feed (and total no. mosquitoes dissected for oocysts, and sporozoites) | Presence of oocysts (and maximum prevalence on a given day) | Presence of sporozoites (and maximum prevalence on a given day) | Day sporozoites first detected in salivary glands | EIP (days) predicted by degree-day model |
| --- | --- | --- | --- | --- | --- | --- |
| *An. stephensi* |  |  |  |  |  |  |
| 16 ± 0 | 3 | 40-62 (50, 92) | No | No | NA | Infinite |
| 17 ± 0 | 2 | 34-60 (64, 273) | Yes (55%) | Yes (57%) | 38 | 111 |
| 18 ± 0 | 3 | 16-62 (402, 322) | Yes (75%) | Yes (73%) | 33 | 56 |
| 20 ± 0 | 1 | 11-39 (148, 114) | Yes (80%) | Yes (78%) | 26 | 28 |
| 14 ± 5 | 3 | 17-60 (144, 217) | Yes (3%) | No | NA | Infinite |
| 16 ± 5 | 6 | 16-55 (362, 377) | Yes (5%) | Yes (5%) | 30 | Infinite |
| 18 ± 5 | 2 | 15-62 (286, 228) | Yes (79%) | Yes (85%) | 27 | 56 |
| *An. gambiae* |  |  |  |  |  |  |
| 17 ± 0 | 2 | 34-50 (107, 273) | Yes (13%) | Yes (3%) | 43 | 111 |
| 19 ± 0 | 2 | 23-41 (52, 383) | Yes (40%) | Yes (23%) | 29 | 37 |
| 19 ± 5 | 2 | 18-40 (80, 491) | Yes (50%) | Yes (33%) | 25 | 37 |

We found no evidence for oocyst or sporozoite infection at constant 16°C over three replicate feeds. Oocyst prevalence in mosquitoes maintained at 27°C (controls) ranged from 82-100%, indicating that these feeds were infectious (Table S1). Positive salivary gland infections were detected at constant 17, 18, and 20°C for *An. stephensi*, and 17 and 19°C for *An. gambiae*. Rates of parasite development were greater than predicted by the established degree-day model.

Adding realistic diurnal temperature fluctuation enabled parasites to establish to oocyst stage at 14 and 16°C, and to complete development to sporozoite stage at 16°C in *An. stephensi*, albeit at low levels (Table 1). In the 18°C and 19°C treatments, temperature variation further shortened EIP for both vector species (Table 1).

Our relatively coarse sampling frequency does not enable us to precisely define EIP (i.e. to definitively capture the day the first individual mosquito becomes infectious). However, by estimating 95% confidence intervals across the replicate infectious feeds we can determine a credible window for EIP, representing the latest day at which no sporozoites were likely to be observed, and the latest day at which maximum sporozoite prevalence was likely to have occurred based on our observations. Our conservative methodology minimized any differences between the observed values and the Detinova values. This approach conceptually follows recent work defining the completion of sporogony as a distribution rather than a single time point [7,15,19]. Using this method we define a window of sporogony for *An. stephensi* of 31-37 days at 18°C, and 26-27 days at 18 ± 5°C (Figure 1A&B). These empirical estimates are substantially shorter than the EIP of 56 days predicted by the degree-day model. The significance of these shorter EIPs can be illustrated by integrating EIP with the respective mosquito survival (Figure 1A&B). The shaded areas provide a measure of the daily number of infectious mosquitoes alive, or ‘infectious-mosquito-days’, which all else being equal, scales with force of infection [15]. Our empirical data suggest a 2.7-fold increase in force of infection at constant 18°C compared with the degree-day model, and 8.5-fold increase at 18 ± 5°C.

**Figure 1.**
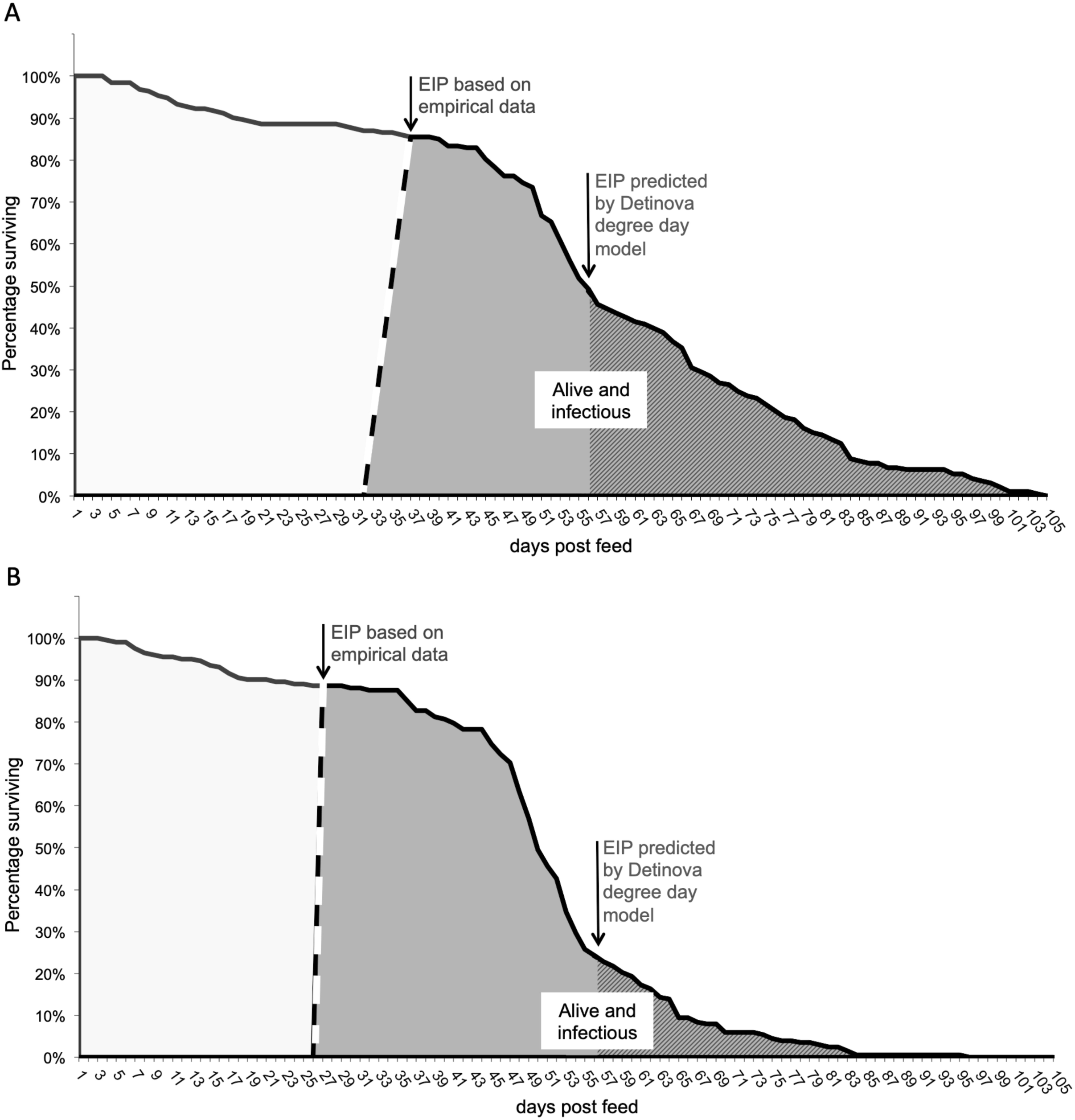
Plot line shows survival of mosquitoes that we assume with become infectious following a parasite-infected blood meal. Areas under the line represent total mosquito days of life, based on empirical data for *An. stephensi* at (A) 18°C, and (B) 18 ± 5°C. The dashed line bounding the grey area represents estimated EIP, or parasite sporogony, based on observed data at these temperatures. The grey area represents the number of infectious-mosquito-days, which provides a relative measure of force of infection. Within the larger grey area, the hatched shading represents infectious-mosquito-days calculated using the degree-day model of Detinova [8].

## 4. Discussion

The Detinova model of EIP has been applied extensively for over 50 years, but with little critical evaluation. Our data suggest that 16°C, the minimum temperature threshold for parasite development assumed in the model, is a good approximation. Adding realistic diurnal temperature fluctuation at 16°C facilitated infection but only 2/377 mosquitoes were positive for sporozoites, so the effect is marginal. A threshold of 16°C is in line with very early empirical work and parasite kinetics research [20,21], but differs from a number of modeling studies that assume a lower limit of 18°C (reviewed in [22]).

However, the Detinova model dramatically underestimates parasite development rate at temperatures marginally above the lower threshold, and fails to capture the effects of realistic daily temperature variation, which enhances development still further. The Detinova model assumes that 111 cumulative degree-days are required to complete sporogony once temperature is above the lower threshold. The current study indicates that the required number of degree-days is not a constant 111 days, but varies non-linearly with temperature. Based on our observed approximations of EIP (Table 1) and the established lower developmental threshold of 16°C, we use the degree-day equation (Equation 1) to calculate that sporogony takes as few as 38, 66 and 104 degree-days at 17, 18 and 20°C, respectively, for *An. stephensi*. For *An. gambiae* we estimate 43 and 87 degree-days at 17 and 19°C, respectively. Note that these estimates do not mean that EIP is shorter in absolute terms at cooler temperatures, but that at temperatures below 21°C it takes proportionally fewer degree-days for parasites to complete sporogony. Moreover, we find that realistic temperature fluctuation can enhance parasite development yet further. These findings are consistent with Jensen’s inequality, which predicts that fluctuation around a mean temperature can modify (in this case increase) development rate compared to the same average constant temperature [23], and confirm earlier empirical research conducted using species of rodent malaria [14]. In turn, the faster parasite development yields potentially much greater force of infection under cooler environmental conditions than predicted using the Detinova model (illustrated in Fig 1).

We acknowledge that our study used lab-adapted mosquito and parasite strains and there is a need to validate our findings using local mosquito-parasite pairings. Local adaptation in vector and/or parasite population could change the thermal sensitivity of the vector-parasite interaction [24]. Note, however, this argument applies equally to the existing Detinova model (and indeed extends to many other mosquito-parasite studies that use lab strains). Similarly, our lab-based mosquito survival curves do not necessarily reflect patterns of survival in the field [25,26], yet qualitative differences in force of infection relative to the Detinova model should hold regardless. As such, our results challenge one of the most long-standing models in malaria biology and highlight a need for further studies to examine the thermal ecology of malaria, particularly at the edges of range in areas such as the Kenyan and Ethiopian highlands where the potential impacts of climate change remain controversial [4,5,27,28].

## Ethical statement

All experiments were conducted under Penn State IBC protocol #48219.

## Data availability

Data from this study are available as electronic supplementary material in Table S1 for all temperatures for mosquitoes dissected for oocysts and sporozoite dissections, sampling intervals, replicates, and sample sizes.

## Author contributions

JLW and ES conducted the experiments. PAL assisted in analysis. JLW and MBT wrote the manuscript with inputs from ES and PAL.

## Competing interests

The authors have no competing interests.

## Funding

This study was part supported by NIH NIAID grant # R01AI110793, NSF grant # DEB-151868, and the USDA NIFA and Hatch Appropriations under Project #PEN04691 and Accession #1018545. The funders had no role in study design, data collection and analysis, decision to publish, or preparation of the manuscript.

### Acknowledgments

We thank MR4/BEI resources for provision of parasites. Thanks to Deonna Soergel, Dean Taylor and Janet Teeple for assistance in the lab, and Anna Guschin and Elizabeth Eswaskio for translating journal articles from Russian to English.

## Supplementary Information

Electronic supplementary material is available online at (link that will be assigned to Table S1 https://…)

